# Nandrolone supplementation promotes AMPK activation and divergent ^18^[FDG] PET brain connectivity in adult and aged mice

**DOI:** 10.1101/2021.10.19.465014

**Authors:** N.S. Strogulski, A Kopczinski, V.G. De Oliveira, R.B. Carteri, G Hansel, G.T. Venturin, S Greggio, J.C. DaCosta, M.A. De Bastiani, M.S. Rodolphi, L.V. Portela

## Abstract

Decreased anabolic androgen levels are followed by impaired brain energy support and sensing with loss of neural connectivity during physiological aging, providing a neurobiological basis for hormone supplementation. Here, we investigated whether nandrolone decanoate (ND) administration mediates hypothalamic AMPK activation and glucose metabolism, thus affecting metabolic connectivity in brain areas of adult and aged mice. Metabolic interconnected brain areas of rodents can be detected by positron emission tomography using ^18^FDG-mPET. Albino CF1 mice at 3 and 18 months of age were separated into 4 groups that received daily subcutaneous injections of either ND (15 mg/kg) or vehicle for 15 days. At the *in vivo* baseline and on the 14^th^ day, ^18^FDG-mPET analyses were performed. Hypothalamic pAMPK^T172^/AMPK protein levels were assessed, and basal mitochondrial respiratory states were evaluated in synaptosomes. A metabolic connectivity network between brain areas was estimated based on ^18^FDG uptake. We found that ND increased the pAMPK^T172^/AMPK ratio in both adult and aged mice but induced ^18^FDG uptake and mitochondrial basal respiration only in adult mice. Furthermore, ND triggered rearrangement in the metabolic connectivity of adult mice and aged mice compared to age-matched controls. Altogether, our findings suggest that ND promotes hypothalamic AMPK activation, and distinct glucose metabolism and metabolic connectivity rearrangements in the brains of adult and aged mice.

## INTRODUCTION

The aging process is associated with a progressive decrease in physiological function, including abnormal nutrient sensing and cerebral hypometabolism. An exacerbation of these biochemical features may evolve to compromised synaptic integrity and connectivity, leading to neurofunctional deficits [1].

In particular, the hypothalamus plays a unique role in integrating brain and whole-body energy metabolism. Hypothalamic cells are sensitive to systemic metabolic signals, such as glucose, fatty acids, and hormones, including, insulin and leptin [2, 3]. Intracellularly, a molecular effector of these energy signals and endocrine regulation is AMP-dependent kinase (AMPK). Once activated, AMPK promotes increased glucose uptake, as well as induction of catabolic pathways leading to ATP production [4–6]. This allows AMPK to dynamically respond to changes in energy demand due to oscillations in synaptic activity, primarily orchestrating substrate flow toward mitochondrial oxidative phosphorylation.

As a metabolic hub connecting different cellular functions, mitochondrial dysfunction associated with aging is believed to contribute to key components leading to cellular senescence, such as exacerbated oxidative stress and promotion of apoptosis [7, 8]. Depending on the localization site at nerve cells, mitochondria can be more vulnerable to damage, exerting a major contribution to aging mechanisms [9]. Based on this, mitochondrial malfunction in synaptic terminals impairs the energy support for neurotransmitter release and uptake and control of ionic homeostasis, which are known substrates for the rupture of synaptic function and neural-cell connectivity [10, 11].

The maintenance of healthy synapses relies on adequate mitochondrial function as well as sufficient metabolic substrate inflow and utilization. Clinically, hallmark neuroimaging findings in aging and age-related disorders include cerebral hypometabolism detected by fluorodeoxyglucose (^18^FDG). Associated with reduced topographical glucose uptake in the brain, diseases common in the elderly population, such as Alzheimer’s and Parkinson’s, display reduced cerebral connectivity assessed by PET-^18^FDG, reflecting neuronal loss and synaptic dysfunction. Interestingly, abnormal energetic homeostasis during aging parallels altered hypothalamic activity [12, 13] and a reduction in hypophysis-hypothalamic-gonadal signaling, leading to reduced testosterone levels in males [14].

Remarkably, many physiological processes in the brain, from development to normal aging, are under the essential influence of the androgenic anabolic steroid (AAS) hormones produced in the gonads. For instance, androgen receptors (ARs) mediate synaptogenesis (synaptic density and dendritic spine number), which is a morphological correlate of neuroplasticity and likely cognitive function [15, 16]. The reported decrease in AAS during the course of aging has been the mechanistic basis for proposing exogenous hormone administration to hamper age-related physiological deficits [17–21]. Testosterone and its synthetic derivative nandrolone decanoate (ND) induce robust anabolic effects in various tissues, including the brain [22–24]. Based on these properties, administration of AAS has already been investigated to attenuate age-related brain pathologies, in which a decrease in bioenergetic function exerts a role in the progression of symptoms and the severity of disease [25–27]. On the other hand, testosterone and ND are often used in young adults to promote anabolic effects such as increased protein synthesis, skeletal muscle mass/strength and oxidative metabolism. However, such an induced metabolic boost is not limited to peripheral tissues and is frequently accompanied by behavioral effects, confirming that beyond morphological effects, AAS also mediates synaptic mechanisms. However, although androgen hormones affect mitochondrial synaptic metabolism and function, the impacts of their use on AMPK-dependent hypothalamic metabolic control and brain connectivity during aging remain elusive.

In this study, we sought to investigate whether ND administration improves ^18^FDG uptake and metabolic connectivity in the brain in adult and aged mice.

## METHODS

### Animal procedures

Albino CF1 mice of 3 and 18 months of age were randomly selected to receive either subcutaneous administration of a synthetic analog of testosterone, nandrolone decanoate (18m/ND) (Decadurabolin^®^, Schering-Plough, Brazil) at a dose of 15 mg/kg, or an equivalent volume of vehicle (Corn oil, Salada^™^, Bunge, Brazil) once a day for 15 consecutive days [24, 28]. Animals were maintained in plexiglass cages, with an average of 4 littermates, and *ad libitum* access to water and standard chow under controlled temperature (22 ±1 °C) and lighting (12 h light/12 h dark, lights on at 7 am) conditions. Following pharmacologic administration, animals were sacrificed, and the hypothalamus was dissected and homogenized for immunoblotting assays. The remaining left cerebral hemisphere was homogenized for synaptosome fractionation, which was used for mitochondrial respirometry. All procedures were performed with the approval of the UFRGS ethics committee for animal experimentation (CEUA UFRGS #33762)

### Immunoblotting analysis

Dissected hypothalamic tissue was homogenized in NP-40 homogenizing buffer, submitted to 12% acrylamide/bis-acrylamide gel electrophoresis (SDS–PAGE) and transferred to and analyzed on nitrocellulose membranes. Membranes were blocked according to each antibody datasheet. Twenty-five micrograms of each sample were incubated overnight at 4°C with pAMPKα2 (Cell Signaling^®^, USA, 1:1000) and AMPKα2 (T172) (Cell Signaling^®^, USA, 1:1000) antibodies, respectively. After overnight incubation, membranes were washed with 1x TTBS and placed in a 1:25000 secondary HRP-conjugated IgG anti-rabbit or anti-mouse solution (GE Healthcare^®^, Sweden) according to the primary antibody datasheet specifications. Western blotting analysis is shown as the percent of control optic density, analyzed using ImageJ software, and normalized to the loading control ß-actin HRP-linked primary antibody (1:10000, Sigma–Aldrich^®^, Saint Louis, MO, USA) according to the manufacturer’s instructions.

### Synaptosome preparation

Animals’ left hemispheres were rapidly dissected and homogenized in 1000 μL of isolation buffer (320 mM sucrose, 10 mM Tris, 0,1% BSA, pH 7.4) using a 10 mL glass potter. The brain tissue was gently homogenized with 10 strokes. The homogenate was immediately used for respirometry and protein quantification for subsequent normalization of oxygen consumption per mg of tissue.

For synaptosome isolation, homogenized samples were then centrifuged at 1330 ×g for 3 min. The supernatant was carefully retained and then centrifuged at 21,200 ×g for 10 min. The resulting pellet was resuspended, carefully layered on top of a discontinuous Percoll gradient, and centrifuged for 5 min at 30,700 ×g [11, 12]. This results in a synaptosome-enriched band that was transferred and resuspended in isolation buffer and centrifuged for an additional 16,900x g for 10 min. The resulting pellet was resuspended in 2 mL isolation buffer and centrifuged at 6,500x g for 10 min. The final pellet of the synaptosome fraction was resuspended in sucrose and Tris buffer (320 mM sucrose, 10 mM Tris, pH 7.4) and immediately used for further analysis as previously described [29, 30].

### MicroPET-scan analysis

Two to six hours before the first subcutaneous injection of ND or VEH, mice were submitted to baseline [^18^F]FDG uptake evaluation using positron emission tomography scans (microPET), performed at the Brain Institute (Porto Alegre, Rio Grande do Sul, Brazil). The same animals were resubmitted to [^18^F]FDG uptake scans again 15 days later. Before both paired scans, animals were fasted overnight but had *ad libitum* access to water. Animals were placed in a controlled temperature environment, and an intraperitoneal injection of 200 µL of [^18^F]FDG with a radioactive activity of approximately 240 μCi was administered, allowing conscious drug uptake for 40 min. Animals were then anesthetized (ketamine/xylazine 90 mg/kg and 7.5 mg/kg, respectively), and microPET scans were performed [31, 32]. Acquisition of ^18^FDG-PET images was mapped and quantified as SUV according to *mirrione* [33], and is expressed as the variation between paired baseline and 15^th^-day scans. Glucose uptake in the brain was topographically assessed in the following regions: (**RSTR; LSTR**: striatum, right and left, respectively.) (**CTX:** cortex) (**RHIP; LHIP:** hippocampus, right and left, respectively) (**THA:** thalamus) (**CB:** cerebellum) (**BFS:** basal forebrain & septum) (**HYP:** hypothalamus) (**RAMY; LAMY**: amygdala, right and left, respectively) (**BS:** brainstem) (**CG:** cingulate gyrus) (**SC**: superior colliculi) (**OLF:** olfactory bulb) (**RMID; LMID:** midbrain, right and left, respectively) (**RIC; LIC:** inferior colliculi, right and left, respectively).

### High-resolution respirometry analysis

High-resolution respirometry was performed using an O2k Oxygraph (Oroboros Instruments, Innsbruck, Austria) utilizing a SUIT (substrate, uncoupler inhibitor titration) protocol to assess specific mitochondrial respiratory states as described by Chance *et al* [34] and redefined by Gnaiger *et al* [35]. Synaptosomes obtained from the left cerebral hemisphere were submitted to a high-resolution respirometric analysis using an O2k oxygraph (Oroboros Instruments, Innsbruck, Austria). The analysis was performed in 300 µL of synaptosomal fraction suspended in 2 mL of respiration buffer (KCI 100 mM, mannitol 75 mM, sucrose 25 mM, KH_2_PO_4_ 5 mM, EDTA 0.05 mM, and Tris-HCl 10 mM, pH 7.4).

The ROUTINE respiration state was reached after the stabilization of oxygen consumption rates following sample addition and is sustained by endogenous energy substrates in which glucose represents a major contribution. After the ROUTINE state was achieved, KCN (1 mM) was added to inhibit complex IV, which limits oxygen consumption to nonoxidative reactions, allowing estimation of ROX. Basal respiration was then estimated as the difference between ROX and ROUTINE states.

### Statistical analysis

Due to the nature of the presented data, an unpaired Student’s t test was performed to assess differences, with significance considered when p<0.05, and normality was investigated using the Kolgomorov-Smirnov normality statistical test. All results in the figures are shown as the mean ± SD. Pearson tests were performed to investigate correlations between variables using the R package *psych* [36], with Bonferroni correction for multiple correlations. Correlation data are expressed as Pearson’s r, and only correlations with r > 0.9 and p<0.01 are shown.

Metabolic interconnected brain areas in rodents can be detected by positron emission tomography using ^18^FDG-mPET. Given this, we estimated metabolic networks through correlation of the [18F]FDG uptake values among different brain anatomical regions of interest (ROIs). The magnitude of the correlation implies the existence of a functional association. For this approach, the Pearson’s r correlation of SUVs obtained in each anatomical region by microPET analysis was used.

Networks were built using the *RedeR* package [37]. Furthermore, we performed clustering with Euclidean distance and the complete linkage agglomeration method using the *pvclust* package [38]; 10000 bootstraps and 123 seed parameters were set to assess the uncertainty in hierarchical clustering. Additionally, heatmaps were obtained from the correlation plots.

## RESULTS

### Nandrolone upregulates hypothalamic AMPK in adult and aged mice

Immunocontent analysis of hypothalamic pAMPK (Thr172) and AMPK in adult and aged mice revealed that ND significantly increased the pAMPK (Thr172)/AMPK ratio in adult mice (3m/ND vs. 3m/ND, p = 0.0274; t = 2.628, df = 9), a proxy indicator of increased AMPK activation (Figure 1A, C). This regulatory effect of nandrolone was also observed in aged mice, with the hypothalamic pAMPK(Thr172)/AMPK ratio significantly increased in 18m/ND mice compared to 18m/VEH (18m/ND vs. vs 18m/VEH, p = 0.0430, t = 2.403, df = 8) (Figure 1B, C).

**Figure 1:**
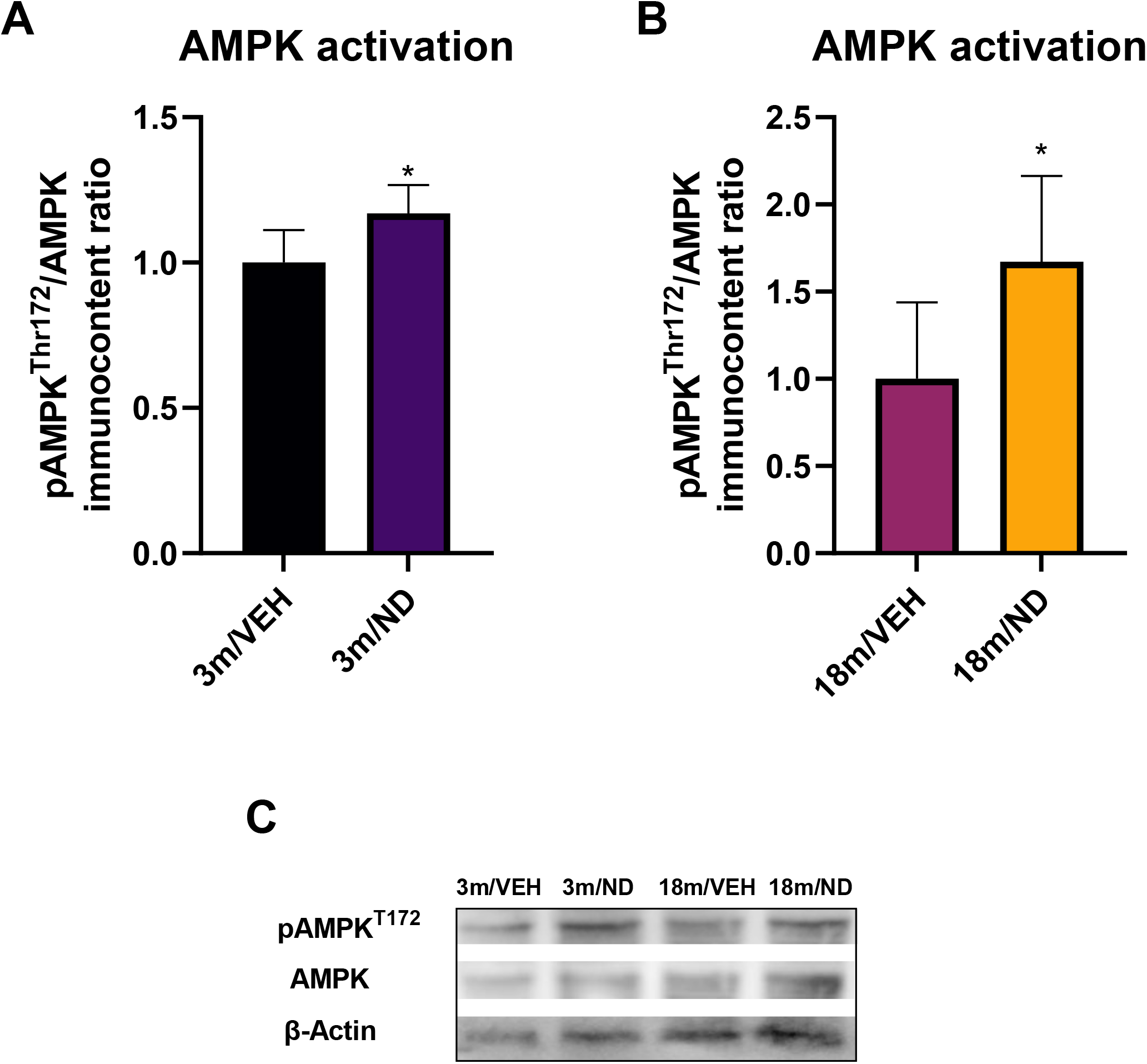
AMPK activation in the hypothalamus. **(A)** pAMPK(T172)/AMPK immunocontent ratio was increased in adult mice relative to age-matched controls (3m/VEH). **(B)** pAMPK(T172)/AMPK immunocontent ratio is increased in aged mice relative to age matched controls (18m/VEH) **(C)** Representative western blotting membrane of AMPK, pAMPK(T172) and loading control β-actin. (Unpaired Student’s t test, * p<0.05; 3m/VEH n = 6, 3m/ND n = 5, 18m/VEH n = 5, 18m/ND n = 5) Results are shown as the mean ± SD.

### Nandrolone supplementation affects glucose metabolism and basal mitochondrial respiration

Brain ^18^FDG uptake after 15 days of nandrolone administration was significantly increased up to 1.5-fold in adult mice (3m/ND group) compared to age-matched controls injected with vehicle (3m/VEH) in all assessed areas (Table 1, and Figure 2A). In contrast, this increase in brain glucose metabolism promoted by ND was not observed in aged mice (18m/ND), compared to aged mice injected with vehicle (18m/VEH) in any of the assessed regions (Table 1 and Figure 2A).

**Table 1:** Descriptive statistics of the alterations promoted by nandrolone in ^18^FDG uptake by brain regions in adult and aged mice. The variation of the baseline and 14 days of nandrolone (ND) or vehicle (VEH) treatment standardized uptake values (ΔSUV) is expressed as mean ± SD. respectively. * indicates p-values <0.05, ** indicates p-values <0.01 and *** indicates p-values < 0.001 (Unpaired Student’s T-test). Abbreviations: df: degrees of freedom, (**RSTR; LSTR**: Striatum, right and left respectively.) (**CTX:** Cortex) (**RHIP; LHIP:** Hippocampus, right and left respectively) (**THA:** Thalamus) (**CB:** Cerebellum) (**BFS:** Basal Forebrain & Septum) (**HYP:** Hypothalamus) (**RAMY; LAMY**: Amygdala, right and left respectively) (**BS:** Brainstem) (**CG:** Cingulate Gyrus) (**SC**: Superior Colliculi) (**OLF:** Olfactory bulb) (**RMID; LMID:** Midbrain, right and left respectively) (**RIC; LIC:** Inferior Colliculi, right and left respectively).

| Region | Average ΔSUV | | df | T-value | p-value |
| --- | --- | --- | --- | --- | --- |
|  | 3m/VEH | 3m/ND |  |  |  |
| **RSTR** | -0.2691 ± 0.66 | 1.4821 ± 0.52 | 9 | -4.8070 | 0.0010 |
| **LSTR** | -0.2498 ± 0.65 | 1.4906 ± 0.50 | 9 | -4.8728 | 0.0009 |
| **CTX** | -0.1957 ± 0.59 | 1.1387 ± 0.34 | 9 | -4.4713 | 0.0016 |
| **RHIP** | -0.1910 ± 0.65 | 1.4296 ± 0.43 | 9 | -4.7440 | 0.0011 |
| **LHIP** | -0.2768 ± 0.60 | 1.4302 ± 0.45 | 9 | -5.2516 | 0.0005 |
| **THA** | -0.2403 ± 0.69 | 1.5623 ± 0.44 | 9 | -5.0065 | 0.0007 |
| **CB** | -0.1233 ± 0.55 | 1.0318 ± 0.35 | 9 | -4.0536 | 0.0029 |
| **BFS** | -0.1619 ± 0.57 | 1.1068 ± 0.45 | 9 | -4.0499 | 0.0029 |
| **HYP** | -0.1415 ± 0.48 | 1.0046 ± 0.39 | 9 | -4.2859 | 0.0020 |
| **RAMY** | -0.0860 ± 0.48 | 0.9737 ± 0.34 | 9 | -4.0947 | 0.0027 |
| **LAMY** | -0.1494 ± 0.50 | 1.0082 ± 0.35 | 9 | -4.3729 | 0.0018 |
| **BS** | -0.1149 ± 0.52 | 1.1135 ± 0.32 | 9 | -4.5572 | 0.0014 |
| **CG** | -0.2406 ± 0.76 | 1.4737 ± 0.42 | 9 | -4.4588 | 0.0016 |
| **SC** | -0.2786 ± 0.72 | 1.4273 ± 0.47 | 9 | -4.5196 | 0.0014 |
| **OLF** | -0.1356 ± 0.62 | 1.2931 ± 0.35 | 9 | -4.5362 | 0.0014 |
| **RMID** | -0.1805 ± 0.70 | 1.4249 ± 0.40 | 9 | -4.5122 | 0.0015 |
| **LMID** | -0.2015 ± 0.63 | 1.4667 ± 0.47 | 9 | -4.8982 | 0.0008 |
| **LIC** | -0.2333 ± 0.74 | 1.5073 ± 0.36 | 9 | -4.7574 | 0.0010 |
| **RIC** | -0.2871 ± 0.67 | 1.2099 ± 0.54 | 9 | -4.0392 | 0.0029 |
|  | 18m/VEH | 18m/ND |  |  |  |
| **RSTR** | -0.0851 ± 0.66 | -0.1602 ± 0.52 | 8 | 0.1458 | 0.8877 |
| **LSTR** | -0.1071 ± 0.63 | -0.1293 ± 0.99 | 8 | 0.0423 | 0.9673 |
| **CTX** | -0.0315 ± 0.53 | -0.1794 ± 0.79 | 8 | 0.3485 | 0.7365 |
| **RHIP** | -0.0072 ± 0.65 | 0.0122 ± 0.99 | 8 | -0.0366 | 0.9717 |
| **LHIP** | 0.0010 ± 0.63 | -0.0062 ± 1.04 | 8 | 0.0133 | 0.9897 |
| **THA** | -0.0253 ± 0.66 | -0.0374 ± 1.05 | 8 | 0.0219 | 0.9831 |
| **CB** | -0.0137 ± 0.61 | -0.1460 ± 0.89 | 8 | 0.2748 | 0.7904 |
| **BFS** | -0.0392 ± 0.53 | -0.1178 ± 0.94 | 8 | 0.1637 | 0.8740 |
| **HYP** | -0.0298 ± 0.50 | 0.0417 ± 0.83 | 8 | -0.1656 | 0.8726 |
| **RAMY** | 0.0110 ± 0.46 | -0.1009 ± 0.75 | 8 | 0.2838 | 0.7838 |
| **LAMY** | -0.0969 ± 0.45 | -0.0157 ± 0.79 | 8 | -0.1994 | 0.8469 |
| **BS** | -0.0097 ± 0.56 | 0.0063 ± 0.89 | 8 | -0.0339 | 0.9738 |
| **CG** | 0.0007 ± 0.76 | 0.0691 ± 1.14 | 8 | -0.1118 | 0.9137 |
| **SC** | -0.0363 ± 0.73 | 0.0069 ± 1.06 | 8 | -0.0752 | 0.9419 |
| **OLF** | -0.0096 ± 0.62 | -0.3017 ± 1.05 | 8 | 0.5339 | 0.6080 |
| **RMID** | 0.0447 ± 0.67 | 0.0323 ± 0.97 | 8 | 0.0235 | 0.9818 |
| **LMID** | 0.0007 ± 0.65 | 0.0122 ± 1.06 | 8 | -0.0206 | 0.9841 |
| **LIC** | -0.0182 ± 0.82 | 0.0281 ± 1.09 | 8 | -0.0758 | 0.9414 |
| **RIC** | -0.0313 ± 0.68 | -0.1148 ± 0.94 | 8 | 0.1610 | 0.8761 |

**Figure 2:**
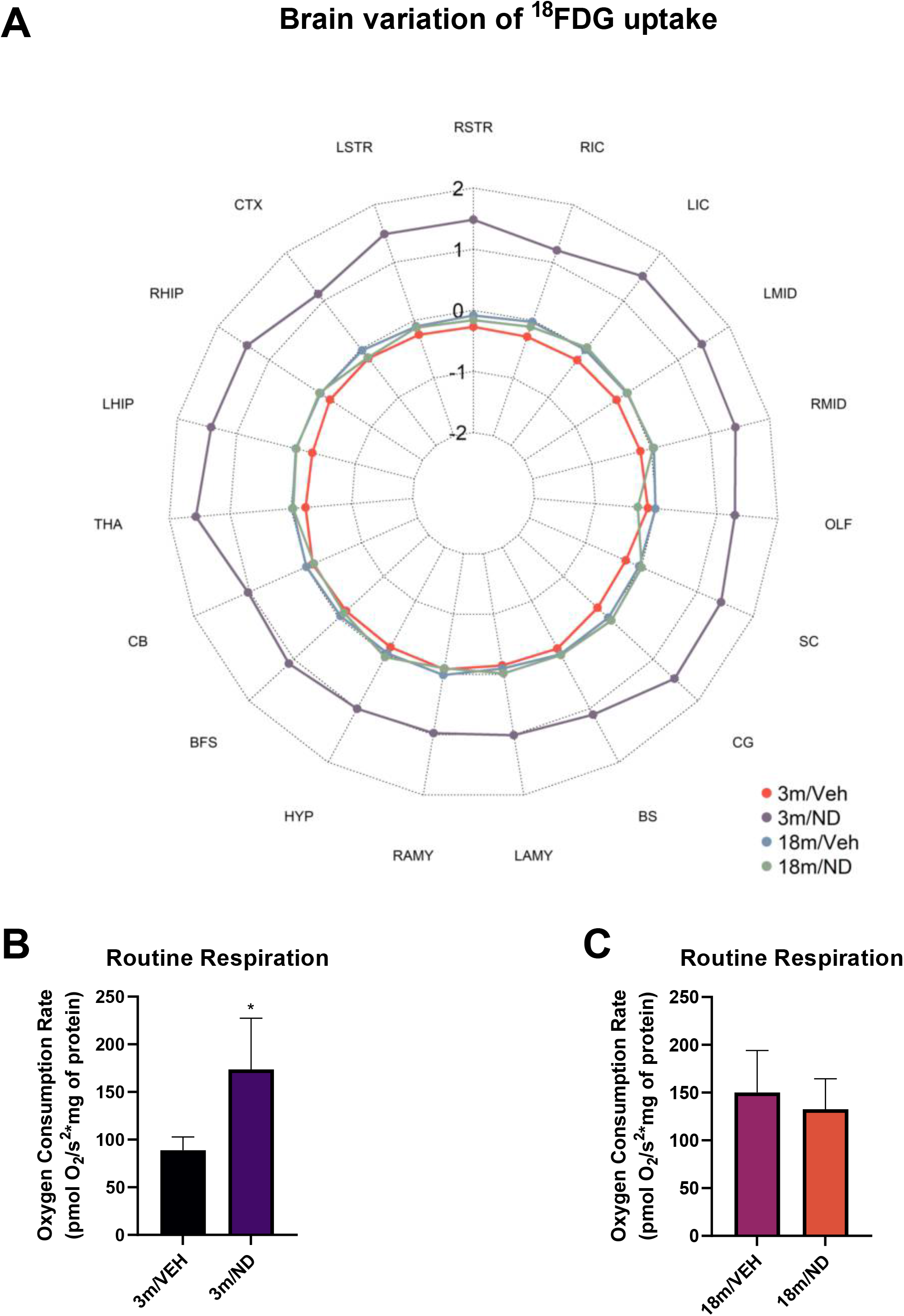
Mitochondrial respiration and ^18^FDG uptake. **(A)** ^18^FDG uptake values of cerebral structures, showing increased uptake in young animals exposed to ND supplementation compared to all groups. Results are shown as the mean variation from baseline ^18^FDG uptake. (3m/VEH n = 6, 3m/ND n = 5, 18m/VEH n = 5, 18m/ND n = 5) **(B)**The routine oxygen consumption rate is increased in adult nandrolone injected mice (3m/ND) compared to aged-matched controls (3m/VEH) (Unpaired Student’s t test, *p = 0.0046, t = 3.748, df = 9). **(C)** Nandrolone did not induce increased basal oxygen consumption rates in synaptic mitochondria of aged mice (18m/ND) compared to age-matched controls (18m/VEH) (Unpaired Student’s t test, p = 0.4899, t = 0.7237, df = 8). Results are shown as the mean ± SD. (3m/VEH n = 6, 3m/ND n = 5, 18m/VEH n = 5, 18m/ND n = 5)

Coupled with ^18^FDG analysis, synaptosomes were submitted to oxygraphic analysis to evaluate the oxidation of endogenous substrates, including glucose. Mitochondria from synaptosome preparations displayed increased oxygen consumption in adult mice supplemented with ND (3m/ND), compared to controls (3m/VEH), (Unpaired Student’s t test, 3m/VEH vs. 3m/ND, p=0.0046, t=3.748, df=9) (Figure 2B). Aged mice supplemented with nandrolone (18m/ND) displayed no significant difference in routine respiration compared to age-matched controls (18 m/VEH) (unpaired Student’s t test, 18m/VEH vs. 18m/ND, p=0.4899, t=0.7237, df=8) (Figure 2C).

A neural connectivity network using the between-region correlation of ^18^PET-FDG is shown in Figure 3. In this network, brain regions were clustered by correlation similarity into containers of closely related areas. The edge width between these containers is associated with connectivity among containers and their composing elements (regions). From this, we observed that ND administration in adult mice (3m/ND) promotes an overall loss of associations, which is visually evidenced by fewer interconnected regions (nodes), fewer intercorrelations inside the containers (links), and finally, the loss of entire correlation clusters relative to age-matched controls (edge width) (3m/VEH) (Figure 3A-B). In contrast, ND supplementation in aged mice (18m/ND) did not elicit an expressive loss in the number of interconnected regions or in the degree of integration in the correlation network, indicated by the number of node and cluster links and edge width, compared to age-matched controls (18m/VEH).

**Figure 3:**
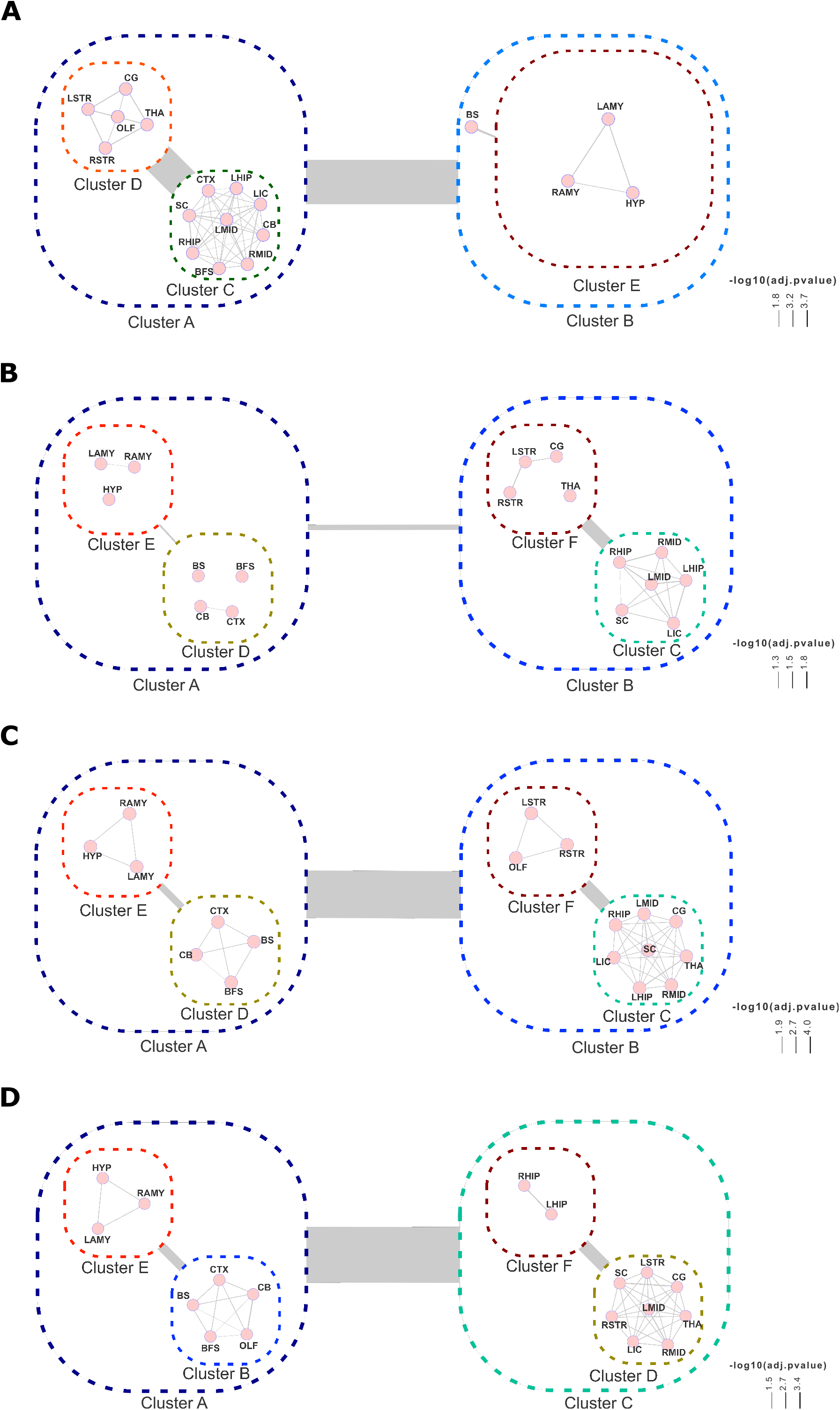
Nandrolone triggers metabolic connectivity rearrangements in the brains of adult and aged mice. **(A, B)** The correlation network of control adult mice (A) was drastically disrupted by 15 days of supplementation with nandrolone (B), exhibiting marked loss of connections between nodes and clusters, indicating compromised synchronous cerebral activity due to supplementation. **(C, D)** Nandrolone supplementation in aged mice alters connectivity, shifts connections, and possibly alters the hierarchy of cerebral interactions but does not lead to reduced cluster or node associations, indicating remodeling but not the overall loss of connections.

## Discussion

Healthy aging in humans is associated with a disruption in brain metabolism and connectivity and a reduction in androgen hormone levels. In this study, we found that age exerts influence over the ND-promoted metabolic rewiring in the brain but was not associated with AMPK activation.

Hormone replacement therapies are employed in aged populations for therapeutic purposes and target age-related declines in physiological function. Additionally, hormones such as ND are frequently used for the treatment of catabolic disorders and by young adults in drug abuse for aesthetic and performance purposes. The metabolic effects of the therapeutic and nontherapeutic use of these drugs on the brains of adult and aged populations are widely unknown.

Here, we demonstrated that ND supplementation induces increased AMPK phosphorylation, a key metabolic regulator in the hypothalamic brain area. This increase in AMPK activation (pAMPK^Thr172^) following the acute use of steroid hormones has been described in other tissues, such as human cardiomyocytes [39], skeletal muscle[40], and the hypothalamic arcuate nucleus of guinea pigs [41]. Herein, we expanded this concept by demonstrating that AMPK activation is not restricted to the arcuate nucleus subregion but may be part of the entire hypothalamic signaling strategy to regulate energy homeostasis. Interestingly, such effects appear to be sustained by ND over the course of at least 15 days in both adult and aged mice. This modulatory effect of steroid hormones may contribute to attenuating certain hallmark features of aging, such as abnormal nutrient sensing, which leads to imbalances in food intake and energy expenditure control in aged individuals [1]. In guinea pigs, activation of the arcuate nucleus AMPK by testosterone treatment led to not only adaptations in the orexigenic neuronal network and feeding patterns but also systemic metabolism, increasing body temperature up to 16 hours post injection through AMPK-dependent mechanisms [41].

The cellular effect of AMPK activation is the promotion of major catabolic pathways, generating endogenous substrates that culminate in increased mitochondrial respiration coupled with ATP synthesis [42, 43]. We showed that ND induces increased mitochondrial oxidation of endogenous substrates only in adult mouse synaptic terminals. Steroid hormones are known to modulate synaptic mitochondrial activity through the translocation of cytoplasmic androgen receptors to nuclei and by directly binding to mitochondrial androgen receptors [44, 45]. Studies have suggested that synaptic mitochondria in ND-treated mice display increased basal mitochondrial membrane potential and hydrogen peroxide production, a proxy indicator of increased endogenous substrate availability and utilization [22], a metabolic feature reinforced by our findings. However, an increase in basal mitochondrial respiration was not observed in ND-supplemented aged mice, indicating a certain level of resistance to this hormone, as evidenced by the limited metabolic responsiveness. A previous work published by Holloway *et al* [46] suggests that metabolic alterations observed in the skeletal muscle of aged males do not translate directly into mitochondrial respirometric alterations due to an increased insensitivity to ADP, leading to increased ROS production throughout the lifespan. We corroborate this notion by showing that although ND supplementation promotes increased AMPK activation, this does not translate to equivalent elevation in basal mitochondrial respiration in aged mice.

Hypothalamic AMPK regulates glucose metabolism in both peripheral organs and the brain. In humans, testosterone has been described to increase ^18^FDG uptake, restoring regional hypometabolism in the brains of both adult men and women [18, 47]. These findings highlight the regulatory effect of androgen hormones in glucose metabolism through increased AMPK activation, which was previously shown in cardiomyocytes [48]. Here, we found that ND supplementation in adult mice increases brain glucose metabolism, further expanding the concept of androgen regulation over hypothalamic control of brain metabolism through AMPK activation.

Intriguingly, ND did not increase brain glucose metabolism in aged mice, which is not coincident with human studies previously reported in the literature. For instance, testosterone replacement therapy (TRT) effects on either cognitively impaired Alzheimer’s disease patients or eugonadal patients have shown increases in cortical glucose metabolism [49, 50]. In our study, the lack of differences between ^18^FDG uptake and basal mitochondrial respiration after ND supplementation may shed light on differences in dose and time regimen influencing metabolic mechanisms in the brains of aged humans and mice. While we used 15 mg/kg ND for 15 days, those authors used doses ranging from 100mg (topical gel, daily) to 200 mg/kg (i.m., quarterly) for up to three months.

Although ND caused an approximately 1.5-fold increase in whole-brain glucose metabolism in adult mice, there was apparent disruption of between-brain area metabolic networks regardless of age, albeit at different intensities. Reductions in brain connectivity patterns in response to androgen steroid administration have been shown using MRI and fMRI neuroimaging techniques. For instance, women injected with testosterone display reduced functional connectivity between the inferior frontal gyrus and the anterior cingulate cortex, which manifested by poorer performance in the ‘Reading the Mind in the Eyes’ test [51, 52]. This suggests that steroid hormone supplementation promotes a loss of connectivity through mechanisms involving alterations in blood flow, substrate uptake, and likely metabolic uncoupling, which may be triggers of synaptic abnormalities and likely neurofunctional deficits.

## Conclusion

In summary, our findings suggest that the increase in AMPK activation in response to ND sustains increased glucose oxidation in the whole brain of adult mice. Overall, brain glucose metabolism was not improved by ND in aged mice despite AMPK activation. These findings suggest that ND promotes a distinct organization in the structure and hierarchy of metabolic connectivity among brain areas in both aged and adult mice.

## Acknowledgments

This work was supported by the Brazilian Agencies/Programs FAPERGS #1010267, FAPERGS/PPSUS#17/2551-0001, FAPERGS/PRONEX#16/2551-0000499-4, Programa de Internacionalização de Ciência FAPERGS/CAPES #19/25510000717-5, Program Science without Borders CNPQ #4011645/2012-6, and CNPq INNT #5465346/2014-6.

## Author Contributions

All authors contributed to the study conception and design. Material preparation, data collection and analysis were performed by Nathan R. Strogulski, Afonso Kopczynski, Vitória Girelli De Oliveira, Randhall Bruce Carteri and Marcelo S. Rodolphi. Micro-PET FDG preparation, processing and analysis were performed by Giannina Venturin, Samuel Greggio and Jaderson C. da Costa. Marco A. de Bastiani performed the brain connectivity analysis. The first draft of the manuscript was written by Nathan R. Strogulski and all authors commented on previous versions of the manuscript. All authors read and approved the final manuscript.

## Data availability

The datasets generated during and/or analysed during the current study are available in the Open Science Framework repository, DOI 10.17605/OSF.IO/K7CX3.

## References

1. López-Otín C, Blasco MA, Partridge L, et al (2013) The Hallmarks of Aging. Cell 153:1194–1217. https://doi.org/http://dx.doi.org/10.1016/j.cell.2013.05.039

2. Andersson U, Filipsson K, Abbott CR, et al (2004) AMP-activated protein kinase plays a role in the control of food intake. The Journal of biological chemistry 279:12005–8. https://doi.org/10.1074/jbc.C300557200

3. Dagon Y, Hur E, Zheng B, et al (2012) p70S6 Kinase Phosphorylates AMPK on Serine 491 to Mediate Leptin’s Effect on Food Intake. Cell Metabolism 16:104–112. https://doi.org/10.1016/j.cmet.2012.05.010

4. Barnes K, Ingram JC, Porras OH, et al (2002) Activation of GLUT1 by metabolic and osmotic stress: potential involvement of AMP-activated protein kinase (AMPK). Journal of cell science 115:2433–42

5. Merrill GF, Kurth EJ, Hardie DG, Winder WW (1997) AICA riboside increases AMP-activated protein kinase, fatty acid oxidation, and glucose uptake in rat muscle. The American journal of physiology 273:E1107–12

6. Habets DDJ, Coumans WA, El Hasnaoui M, et al (2009) Crucial role for LKB1 to AMPKalpha2 axis in the regulation of CD36-mediated long-chain fatty acid uptake into cardiomyocytes. Biochimica et biophysica acta 1791:212–9. https://doi.org/10.1016/j.bbalip.2008.12.009

7. Chapman J, Fielder E, Passos JF (2019) Mitochondrial dysfunction and cell senescence: deciphering a complex relationship. FEBS Letters 593:1566–1579. https://doi.org/10.1002/1873-3468.13498

8. Torres AK, Jara C, Park-Kang HS, et al (2021) Synaptic Mitochondria: An Early Target of Amyloid-β and Tau in Alzheimer’s Disease. Journal of Alzheimer’s Disease 84:1391–1414. https://doi.org/10.3233/JAD-215139

9. Brown MR, Sullivan PG, Geddes JW (2006) Synaptic mitochondria are more susceptible to Ca2+overload than nonsynaptic mitochondria. The Journal of biological chemistry 281:11658–11668. https://doi.org/10.1074/jbc.M510303200

10. Rangaraju V, Calloway N, Ryan TA (2014) Activity-Driven Local ATP Synthesis Is Required for Synaptic Function. Cell 156:825–835. https://doi.org/10.1016/j.cell.2013.12.042

11. Rangaraju V, Lauterbach M, Schuman EM (2019) Spatially Stable Mitochondrial Compartments Fuel Local Translation during Plasticity. Cell 176:73–84.e15. https://doi.org/10.1016/j.cell.2018.12.013

12. Kakimoto A, Ito S, Okada H, et al (2016) Age-Related Sex-Specific Changes in Brain Metabolism and Morphology. 221–226. https://doi.org/10.2967/jnumed.115.166439

13. Brendel M, Focke C, Blume T, et al (2017) Time Courses of Cortical Glucose Metabolism and Microglial Activity Across the Life-Span of Wild-Type Mice: A PET Study. Journal of Nuclear Medicine 49:jnumed.117.195107. https://doi.org/10.2967/jnumed.117.195107

14. Mitchell R, Hollis S, Rothwell C, Robertson WR (1995) Age related changes in the pituitary-testicular axis in normal men; lower serum testosterone results from decreased bioactive LH drive. Clin Endocrinol (Oxf) 42:501–507

15. Jia J, Cui C, Yan X, et al (2016) Effects of testosterone on synaptic plasticity mediated by androgen receptors in male SAMP8 mice. Journal of Toxicology and Environmental Health, Part A 79:. https://doi.org/10.1080/15287394.2016.1193113

16. Devoogd TJ, Nixdorf B, Nottebohm F (1985) Synaptogenesis and changes in synaptic morphology related to acquisition of a new behavior. Brain Research 329:. https://doi.org/10.1016/0006-8993(85)90539-6

17. Bahrke MS, Iip EY, Wright JE (1996) Psychological and Behavioural Effects of Endogenous Testosterone and Anabolic-Androgenic Steroids. 22:367–390

18. Miller KK, Deckersbach T, Rauch SL, et al (2004) Testosterone administration attenuates regional brain hypometabolism in women with anorexia nervosa. 132:197–207. https://doi.org/10.1016/j.pscychresns.2004.09.003

19. Howell SJ, Radford JA, Smets EMA, Shalet SM (2000) Fatigue, sexual function and mood following treatment for haematological malignancy: the impact of mild Leydig cell dysfunction. British Journal of Cancer 82:789–793

20. Burris A, Banks S, Carter C, et al (1992) A Long-term, Prospective Study of the Physiologic and Behavioural Effects of Hormone Replacement in Untreated Hypogonadal Men. Journal of Andrology 13:297–304

21. Brown-Séquard C (1899) Du role physiologique et thérapéutique d’un suc extrait de testicules d’animaux d’après nombre des faix observés chez l’homme. Archives de Physiologie Normale et Pathologique 1:739–746

22. Carteri RB, Kopczynski A, Menegassi LN, et al (2019) Anabolic-androgen steroids effects on bioenergetics responsiveness of synaptic and extrasynaptic mitochondria. Toxicology letters 307:72–80. https://doi.org/10.1016/j.toxlet.2019.03.004

23. Rodolphi MS, Kopczynski A, Carteri RB, et al (2021) Glutamate transporter-1 link astrocytes with heightened aggressive behavior induced by steroid abuse in male CF1 mice. Hormones and Behavior 127:104872. https://doi.org/10.1016/j.yhbeh.2020.104872

24. Kalinine E, Zimmer ER, Zenki KC, et al (2014) Nandrolone-induced aggressive behavior is associated with alterations in extracellular glutamate homeostasis in mice. Hormones and Behavior 66:383–392. https://doi.org/10.1016/j.yhbeh.2014.06.005

25. Carteri RB, Kopczynski A, Rodolphi MS, et al (2019) Testosterone Administration after Traumatic Brain Injury Reduces Mitochondrial Dysfunction and Neurodegeneration. Journal of Neurotrauma 36:2246–2259. https://doi.org/10.1089/neu.2018.6266

26. Singh IN, Sullivan PG, Deng Y, et al (2006) Time Course of Post-Traumatic Mitochondrial Oxidative Damage and Dysfunction in a Mouse Model of Focal Traumatic Brain Injury: Implications for Neuroprotective Therapy. Journal of Cerebral Blood Flow & Metabolism 26:1407–1418. https://doi.org/10.1038/sj.jcbfm.9600297

27. Bianchi VE, Rizzi L, Bresciani E, et al (2020) Androgen Therapy in Neurodegenerative Diseases. Journal of the Endocrine Society 4:. https://doi.org/10.1210/jendso/bvaa120

28. Rodolphi MS, Kopczynski A, Carteri RB, et al (2020) Glutamate transporter-1 link astrocytes with heightened aggressive behavior induced by steroid abuse in male CF1 mice. Hormones and behavior 127:104872. https://doi.org/10.1016/j.yhbeh.2020.104872

29. Sims NR, Anderson MF (2008) Isolation of mitochondria from rat brain using Percoll density gradient centrifugation. Nature Protocols 3:1228–1239. https://doi.org/10.1038/nprot.2008.105

30. Sims NR (1990) Rapid isolation of metabolically active mitochondria from rat brain and subregions using Percoll density gradient centrifugation. Journal of neurochemistry 55:698–707

31. Zanirati G, Azevedo PN, Venturin GT, et al (2018) Depression comorbidity in epileptic rats is related to brain glucose hypometabolism and hypersynchronicity in the metabolic network architecture. Epilepsia. https://doi.org/10.1111/epi.14057

32. Silva RBM, Greggio S, Venturin GT, et al (2018) Beneficial Effects of the Calcium Channel Blocker CTK 01512-2 in a Mouse Model of Multiple Sclerosis. Molecular Neurobiology 1–21. https://doi.org/10.1007/s12035-018-1049-1

33. Mirrione MM, Schiffer WK, Siddiq M, et al (2006) PET imaging of glucose metabolism in a mouse model of temporal lobe epilepsy. Synapse (New York, NY) 59:119–121. https://doi.org/10.1002/syn.20216

34. Chance B, Williams GR (1955) Respiratory enzymes in oxidative phosphorylation. I. Kinetics of oxygen utilization. The Journal of biological chemistry 217:383–93

35. Gnaiger E, Gnaiger E (2012) Mitochondrial Pathways and Respiratory Control An Introduction to OXPHOS Analysis Mitochondrial Pathways and Respiratory Control An Introduction to OXPHOS Analysis

36. Revelle W (2018) psych: Procedures for Psychological, Psychometric, and Personality Research

37. Castro MAA, Wang X, Fletcher MNC, et al (2012) RedeR: R/Bioconductor package for representing modular structures, nested networks and multiple levels of hierarchical associations. Genome biology 13:R29. https://doi.org/10.1186/gb-2012-13-4-r29

38. Suzuki R, Shimodaira H (2006) Pvclust: an R package for assessing the uncertainty in hierarchical clustering. Bioinformatics (Oxford, England) 22:1540–2. https://doi.org/10.1093/bioinformatics/btl117

39. Troncoso MF, Pavez M, Wilson C, et al (2021) Testosterone activates glucose metabolism through AMPK and androgen signaling in cardiomyocyte hypertrophy. Biological research 54:3. https://doi.org/10.1186/s40659-021-00328-4

40. Ghanim H, Dhindsa S, Batra M, et al (2020) Testosterone Increases the Expression and Phosphorylation of AMP Kinase α in Men With Hypogonadism and Type 2 Diabetes. The Journal of clinical endocrinology and metabolism 105:1169–1175. https://doi.org/10.1210/clinem/dgz288

41. Borgquist A, Meza C, Wagner EJ (2015) The role of AMP-activated protein kinase in the androgenic potentiation of cannabinoid-induced changes in energy homeostasis. 482–495. https://doi.org/10.1152/ajpendo.00421.2014

42. Lage R, Dieguez C, Vidal-Puig A, Lopez M (2008) AMPK: a metabolic gauge regulating whole-body energy homeostasis. Trends in molecular medicine 14:539–549. https://doi.org/10.1016/j.molmed.2008.09.007

43. Hardie DG, Ross FA, Hawley SA (2012) AMPK: a nutrient and energy sensor that maintains energy homeostasis. Nature reviews Molecular cell biology 13:251–262. https://doi.org/10.1038/nrm3311

44. Bajpai P, Koc E, Sonpavde G, et al (2019) Mitochondrial localization, import, and mitochondrial function of the androgen receptor. Journal of Biological Chemistry 294:. https://doi.org/10.1074/jbc.RA118.006727

45. Molano F, Saborido A, Delgado J, et al (1999) Rat liver lysosomal and mitochondrial activities are modified by anabolic-androgenic steroids. Medicine & Science in Sports & Exercise 31:243–250. https://doi.org/10.1097/00005768-199902000-00007

46. Holloway GP, Holwerda AM, Miotto PM, et al (2018) Age-Associated Impairments in Mitochondrial ADP Sensitivity Contribute to Redox Stress in Senescent Human Skeletal Muscle. Cell Reports 22:2837–2848. https://doi.org/10.1016/J.CELREP.2018.02.069

47. Zitzmann M, Weckesser M, Schober O, Nieschlag E (2001) Changes in cerebral glucose metabolism and visuospatial capability in hypogonadal males under testosterone substitution therapy. Experimental and Clinical Endocrinology & Diabetes 109:302–304. https://doi.org/10.1055/s-2001-16351

48. Troncoso MF, Pavez M, Wilson C, et al (2021) Testosterone activates glucose metabolism through AMPK and androgen signaling in cardiomyocyte hypertrophy. Biological research 54:3. https://doi.org/10.1186/s40659-021-00328-4

49. Tan RS (2013) Testosterone effect on brain metabolism in elderly patients with Alzheimer’s disease: comparing two cases at different disease stages. Aging clinical and experimental research 25:343–7. https://doi.org/10.1007/s40520-013-0049-2

50. Maki PM, Ernst M, London ED, et al (2007) Intramuscular testosterone treatment in elderly men: evidence of memory decline and altered brain function. The Journal of clinical endocrinology and metabolism 92:4107–14. https://doi.org/10.1210/jc.2006-1805

51. Bos PA, Hofman D, Hermans EJ, et al (2016) Testosterone reduces functional connectivity during the “Reading the Mind in the Eyes” Test. Psychoneuroendocrinology 68:194–201. https://doi.org/10.1016/j.psyneuen.2016.03.006

52. Votinov M, Wagels L, Hoffstaedter F, et al (2020) Effects of exogenous testosterone application on network connectivity within emotion regulation systems. Scientific Reports 10:. https://doi.org/10.1038/s41598-020-59329-0

